# Analysis of phage-resistant mechanisms in *Staphylococcus aureus* SA003 reveals a different binding mechanism for the closely related Twort-like phages ϕSA012 and ϕSA039

**DOI:** 10.1101/339549

**Authors:** Aa Haeruman Azam, Fumiya Hoshiga, Ippei Takeuchi, Kazuhiko Miyanaga, Yasunori Tanji

## Abstract

We have previously generated strains of *Staphylococcus aureus* SA003 resistant to its specific phage ϕSA012 through long-term coevolution experiment. However, the DNA mutations responsible for the phenotypic change of phage resistance are unknown. Whole-genome analysis revealed six genes that acquired unique point mutations: five missense mutations and one nonsense mutation. Moreover, one deletion, 1.779-bp, resulted in the deletion of the genes encoding glycosyltransferase, TarS, and iron-sulfure repair protein, ScdA. The deletion occurred from the second round of coculture (SA003R2) and remained through the last round. The ϕSA012 infection toward SA003R2 had decreased to 79.77±7.50% according to plating efficiency. Complementation of the phage-resistant strain by the wild-type allele showed two mutated host genes were linked to the inhibition of post-adsorption, and five genes were linked to phage adsorption of ϕSA012. Unlike ϕSA012, infection by ϕSA039, a close relative of ϕSA012, onto SA003R2 was impaired drastically. Complementation of SA003R2 by wild-type *tarS* restores the infectivity of ϕSA039. Thus, we concluded that ϕSA039 requires β-GlcNAc in Wall Teichoic Acid (WTA) for its binding. In silico analysis of the ϕSA039 genome revealed that several proteins in the tail and baseplate region were different from ϕSA012; notably the partial deletion of *orf96* of ϕSA039, a homolog of *orf99* of ϕSA012. *Orf100* of ϕSA039, a homolog of *Orf103* of ϕSA012, a previously reported receptor binding protein (RBP), had low similarity (86%) to that of ϕSA012. The difference in tail and baseplate proteins might be the factor for specificity difference between ϕSA012 and ϕSA039.

## Introduction

*Staphylococcus aureus* infection in humans and animals is of worldwide concern due to the emergence of antibiotic-resistant strains such as methicillin-resistant *S. aureus* (MRSA) and vancomycin-resistant *S. aureus* (VRSA) (Enright et al. 2002; Sakoulas et al. 2005). In dairy industry, for instance, *S. aureus* is one of the most frequent causative agent of bovine mastitis, with prevalence rates as high as 50% in some countries (Leitner et al. 2003). Due to its negative effect on milk quality, high quantities of expensive antibiotic are frequently used. Alternatively, using bacteriophage as a means of controlling *S. aureus* is a promising treatment because of the high specificity and lytic activity of the phage. Various efforts are currently made to evaluate the potential of phage therapy (Matsuzaki et al. 2005; Maciejewska et al. 2018). In order to develop phage therapy for mastitis, we isolated several *S. aureus* stains, including *S. aureus* SA003, from the milk of mastitic cows in our previous study (Synnott et al. 2009). We also isolated two lytic phages with wide host ranges, ϕSA012 and ϕSA039, from sewage influent (Synnott et al. 2009). However, in batch co-culture experiment, *S. aureus* developed resistance to ϕSA012 (Synnott et al. 2009). The emergence of phage-resistant bacteria is a big obstacle to realizing phage therapy. A deeper understanding of phage resistance mechanisms is critical before we apply phage therapy in real world.

Phage-resistant bacteria typically evolve phage adsorption inhibition by, for example, altering their phage receptor (Bohannan and Lenski 2000; Denes et al. 2015). Though, there is a report that showed most phage-resistant strains of *Streptococcus thermophilus* used the Clustered Regularly Interspaced Short Polindrom Repeat (CRISPR) system to defend against phage infection (Levin et al. 2013). At the genomic level, mechanisms of phage adsorption inhibition have been well-studied in several Gram-negative bacteria (Morona et al. 1985; Marston et al. 2012; Meyer et al. 2012; Seed et al. 2012; Castillo et al. 2015) and a few Gram-positive bacteria, such as *Listeria monocytogenes* (Denes et al. 2015) and *Streptococcus pneumoniae* (Leprohon et al. 2015). However, phage resistance mechanism of *S. aureus* and the specific genes that contribute to the phage resistance are not well studied.

In response to phage resistance bacteria, phage has ability to counteradapt which is called as coevolution of phage and bacteria (Golais et al. 2013). This phenomenon was also observed in *E. coli* O157:H7 co-cultured with phage PP01 (Mizoguchi et al. 2003; Fischer et al. 2004) and O157:H7 co-cultured with coliphages SP-21, SP-22, and SP-15 (Tanji et al. 2005). Previously, we demonstrated the coevolution of *S. aureus* SA003 and phage ϕSA012 during their long-term co-culture (Osada et al. 2017). We observed cumulative development of phage resistance of the host and corresponding infectivity of the phages during this coevolution (Osada et al. 2017). The coevolution of *S. aureus* and phage is unique and should be taken into account when considering the result of clinical trials. In this paper, we presented the DNA mutation responsible for phenotypic change during coevolution experiment of *S. aureus* SA003 and phage ϕSA012.

The phenotype as a result of coevolutionary adaptation of host and phage has previously been observed and explained in our previous report (Takeuchi et al. 2016; Osada et al. 2017). In this study, we focused on genetic analysis of the host cell by whole genome sequencing (WGS). WGS of wild-type SA003 and its phage resistance derivatives enables us to identify the host genes that contribute to the phage resistance during coevolution. Understanding host genes that endow phage resistance may help us design a better strategy for applying phage therapy under real-world conditions.

For the treatment of *S. aureus* and other *Staphylococcus* species, *Myoviridae* phages are considered to be promising candidates for phage therapy (Loessner et al. 1996; Alves et al. 2014). Phages belonging to this genera have been reported to have the following features: being strictly virulent, genome size of 127 to 140 kbp, share considerable similarity at the protein sequence level, and infecting Gram positive bacteria with low GC content (Łobocka et al. 2012). Our phages ϕSA012 and ϕSA039 (from our collection), which belongs to *Myoviridae* genus Twort-like phage, displayed to have different host preferences and morphology (Synnott et al. 2009). In mice, the application of ϕSA012 has shown an effective and promising outcome (Iwano et al. 2018). However, little is known about why these two closely related phages (ϕSA012 and ϕSA039) have difference host preferences. In this study, we identify the different binding mechanism of these two phages and analyze WGS of the phages to find out the gene that contribute to the host-specificity different. Elucidating the details of the host recognition mechanism of these two different Twort-like phage species will expand our understanding of how closely related phages exhibit different host preferences. Our finding provides substantial information for expanding the utility of staphylococcal Twort-like phages in the future for practical use.

## Materials and Methods

### Bacteria, phages, and plasmids

Bacteria, phages, and plasmids used in this study are listed in **Table 1**. *S. aureus* SA003 was previously isolated from milk of mastitic cow and used for propagation of phages. *S. aureus* strain RN4220 was kindly provided by Prof. Motoyuki Sugai (Hiroshima University Graduate School of Biomedical & Health Science, Hiroshima, Japan) with the permission of Richard P, Novick (Skirball Institute of Biomolecular Medicine, New York, NY) and used for genetic manipulation. The *S. aureus* virulent phage ϕSA012 and ϕSA039 were isolated from sewage in Japan (Synnott et al. 2009). Plasmid pLI50 was purchased from Addgene (Cambridge, MA, USA). Plasmid pLIP3 was constructed using pLI50 and P3 promoter, which is constitutive in *S. aureus* (Lee et al. 1991; Jeong et al. 2011; Takeuchi et al. 2016). Plasmid pKOR1-mcs was kindly provided by Dr. Taeok Bae (Indiana University School of Medicine—Northwest, Indianapolis, IN). Plasmid pCasSA was a gift from Dr. Quanjiang Ji (ShanghaiTech University, School of Physical Science and Technology, Shanghai, China). All bacteria and phages were stored in 15% glycerol at −80°C. Luria Bertani (LB), brain heart infusion broth, and Trypticase soy broth are used as a liquid media. For growth on agar, the medium was solidified by adding 1.5% (w/v) agar. ϕSA012 and *S. aureus* SA003 were deposited in the culture collection of the NITE Biological Research Center, Kisarazu, Japan under accession numbers NBRC110649 and NBRC110650, respectively (Takeuchi et al. 2016). The phage ϕSA039 is only preserved in our laboratory. For contact details and acquire channel, refer to the address of corresponding author.

### Passage co-culture experiment of *S. aureus* SA003 and ϕSA012

Serial passage of ϕSA012 co-cultured with SA003 was carried out in previous study (Takeuchi et al. 2016; Osada et al. 2017). Briefly, SA003 was inoculated into LB medium and cultured until early exponential phage. Phage ϕSA012 was added with multiplicity of infection (MOI) =0.1. After 2 to 10 days, bacterium-phage mixed cultures were transferred into new LB medium (1:100 dilution) and cultured under the same conditions. The co-culture was repeated until 38^th^ passage. A phage-resistant strain of SA003 and mutant ϕSA012 phage were collected at the end of each cycle. The resistant strain was named SA003R**n**, whereas the mutant phage was named ϕSA012M**n**, where **n** represents the passage number (e.g., SA003<u>R11</u> refers to the phage-resistant SA003 derivative isolated from co-culture at the <u>11</u>^th^ passage, while ϕSA01<u>2M11</u> refers to the mutant phage ϕSA012 isolated from the co-culture at the <u>11</u>^th^ passage).

### Molecular cloning in *S. aureus*

The primers used in this study are listed in **Table 2**. The primers were purchased from Hokkaido System Science (Japan) and Integrated DNA Technologies (Singapore). T4 DNA ligase and restriction enzymes were purchased from New England Biolabs (New England BioLabs, Ipswich, MA, USA). Plasmid and PCR product were purified using the Nucleospin^®^ kit (Macherey-Nagel, Duren, Germany). Colony or plaque PCR was performed using DirectAmpPCR^®^ (TAKARA, Shiga, Japan) and processed as recommended by the manufacturer. For PCR using pure DNA, Taq DNA polymerase (TAKARA, Shiga, Japan) was used. Deletions of the genes *S2356* and *S2121*were generated by allelic exchange using plasmid pKOR1-mcs (Bae and Schneewind 2006). Gene disruption in *S497* was performed using the Targetron^®^ gene knockout system (Sigma-Aldrich, St. Louis, MO, USA) (Yao et al. 2006). Deletion of the gene encoding O-acetyltransferase (OatA), *oatA*, and *S156* were performed by CRISPR/Cas9 system using plasmid pCasSA (Chen et al. 2017). The complementation experiment was carried out using *E. coli* / *S. aureus* shuttle vector, pLI50, under P3 promoter (Lee et al. 1991; Jeong et al. 2011). Wild-type alleles of SA003 were used as the insert. Electrocompetent *S. aureus* cells were generated and transformed as previously reported (Monk et al. 2012). To transform SA003R11, a large mass of plasmid (< 20 μg) was first concentrated by Pellet Paint^®^ (Novagen^®^, Billerica, MA, USA) before being introduced into electrocompetent cells. Constructed plasmid was cloned in *E. coli* JM109 and pre-introduced into the *S. aureus* restriction-deficient strain, RN4220, before transforming it into SA003.

### Phage preparation

All phages were propagated by the plate lysate method (Stephenson 2010). Briefly, 100 μl of phage lysate (10^5^ PFU/ml) was mixed with 100 μl of overnight bacterial culture in 3 ml of 0.5% top agar, plated on LB agar, and incubated at 37°C overnight. After 5 ml of salt magnesium (SM) buffer (100 mM NaCl, 8 mM MgSO_4_, 50 mM Tris-HCl [pH 7.5], 0.01% gelatin) was added to the plate and the over layer was scraped off to extract phage, the supernatant was collected by centrifugation (5,000 g, 15 min, 4°C). The obtained phage lysate was purified by polyethylene glycol (PEG) precipitation and CsCl density gradient centrifugation (Stephenson 2010). Each phage culture was titrated and stored at 4°C until used.

### Spot testing

Two microliters of phage lysate at a titer 10^5^ plaque forming units (PFU) was dropped on a lawn of bacteria and incubated overnight at 37°C. Infectivity was determined by assessing the turbidity of plaques in the spots where the phage was dropped.

### Adsorption assay

The adsorption efficiency of phages on *S. aureus* strains was measured by titrating free phage present in the supernatant after defined periods of cell-phage contact. *S. aureus* cells were prepared by 10% inoculation of overnight culture into 4.5 ml of LB medium. The culture was subsequently incubated at 37°C with shaking at 120 rpm to an OD_660_ of 1.0 (10^9^ CFU/ml). Phage lysate (10^7^ PFU/ml) was then added to the bacterial culture. After infection at 37°C with shaking at 120 rpm, free phage was collected by centrifugation (9730 g, 1 min) at defined times and titrated using SA003. For longer incubation, 50 μg/ml of either chloramphenicol or erythromycin was added, and cells were equilibrated for 10 min at 37°C before infection to inhibit cell growth and phage development during incubation with phages (Baptista et al. 2008). Adsorption efficiency was calculated by dividing the number of adsorbed phages by the initial number of phages.

### DNA extraction, sequencing, and bioinformatic

DNA was extracted from bacteria using a DNA extraction kit (Sigma-Aldrich, St. Louis, MO, USA). Whole-genome sequencing of bacteria was conducted with the Illumina Miseq platform. Genomes were assembled by Platanus *de novo* assembly ver1.2.1 (Kajitani et al. 2014). ORFs were predicted and annotated by using the RAST server (http://rast.nmpdr.org/). Genome was mapped by Bowtie2 (Langmead and Salzberg. 2012), and SNPs were detected by SAMtools/BCFtools (Li et al. 2009). Mutated regions were confirmed by PCR and Sanger-sequencing. DNA extraction and genome analysis of phage was performed as previously reported (Takeuchi et al. 2016).

### Measuring phage inhibition by peptidoglycan

Pure peptidoglycan from *S. aureus* (PGN-SA, InvivoGen, Shatin, Hong Kong) was used for our phage inhibition assay. The teichoic acid was removed from peptidoglycan by treatment of 48% hydrofluoric acid at 4°C for 48 hours (De Jonge et al. 1992). Efficiency of plating (EOP) with the addition of peptidoglycan was performed as follows: 10 μl of peptidoglycan solution was added into 100 μl (10^4^ PFU/ml) phage and incubated at room for 1 h. Next, 100 μl of the mixture was mixed with 100 μl *S. aureus* overnight culture in 3 ml of 0.5% top agar, plated on LB agar, and incubated at 37°C overnight.

### Phosphate quantification of WTA and total sugar quantification of whole cell

Wall Teichoic Acid (WTA) was extracted from bacteria culture that has been adjusted to same Colony Forming Unit (CFU). WTA extraction was conducted following previous method (Meredith et al. 2008). Phosphate content of WTA was determined following Chen et al (1956). Same volume of extracted WTA sample and freshly prepared reagent C (mixture of one volume of 6N sulfuric acid with 2 volumes of distilled water, one volume of 2.5% ammonium moybdate, and one volume of 10% ascorbic acid) was mixed, cap with parafilm, and incubated at 37°C for 1.5 – 2 hours, KH_2_PO_4_ was used as phosphate standard. The mixture was cooled down at room temperature for few minutes and read at the absorbance of 820 nm. Total carbohydrate of bacterial cell was measured by Anthrone reagent (Viles and Silverman 1949); 2 ml bacteria culture (2×10^9^CFU) was washed by Phosphate Buffer Saline (PBS) twice and mixed with Anthrone solution (1 mg/ml in concentrated H_2_SO_4_ [95-98% w/v]), the mixture was incubated at 90°C for 10 minutes and the OD_630_ of sample was measured. D(+)-Glucose was used a standard.

### Phage inhibition by antibodies

EOP with antibodies specific for ϕSA012 ORF103 (anti-ORF103) and ORF105 (anti-ORF105) was conducted according to a previous study (Takeuchi et al. 2016).

### Statistical analysis

Two-tailed Student’s *t* test was used to determine statistical significance.

### Accession number(s)

*S. aureus* SA003 and ϕSA039 were submitted to the DDBJ database under the accession number **AP018376** and **AP018375**, respectively. While genome of ϕSA012 was available in the GenBank database under accession number: **AB903967**.

## Results

### Whole genome analysis of phage-resistant SA003

The whole genomes of five phage-resistant mutant strains, as well as parent strain SA003, were sequenced on an Illumina Miseq platform. Mutations detected in the sequenced strains were mapped against the wild-type SA003 reference genome (**Table. 3**). Clustered regularly interspaced polindromic repeat (CRISPR/cas), which is well known as a major genetic barrier against phage infection, is likely absent in the genome of SA003.

A total of eight mutations were identified. One point mutation (AAA→ATA) in *S2356*, which encodes Ferredoxyn glutamate synthase, was found in the phage resistant isolate from the first round, SA003R1. The mutation is preserved until the last round. However, compared to wild-type SA003, the sensitivity of SA003R1 towards ϕSA012 phage did not differ significantly according to plating efficiency and adsorption assays (data not shown). Deleting *S2356* also did not alter the sensitivity of *S. aureus* (data not shown). We assumed that the mutation in *S2356* did not contribute to phage resistance.

The phage-resistant strain in the second round (SA003R2) acquired a point mutation in (CAC→TAC) in *S497* encoding Undecaprenyl-phosphate N-acetylglucosaminyl 1-phosphate transferase, which is well known as teichoic acid ribitol *(tagO)* gene. This strain also acquired an 1779-bp deletion in WTA tar gene cluster gene consisting of: a 1200-bp C-terminal region of *S156* (which encodes glycosyltranferase, TarS); a 125-bp noncoding region; and a 454-bp N-terminal region of *S157* (which encodes iron-sulfur repair protein, Scd) (**Fig. 1A**).

**Figure 1.**
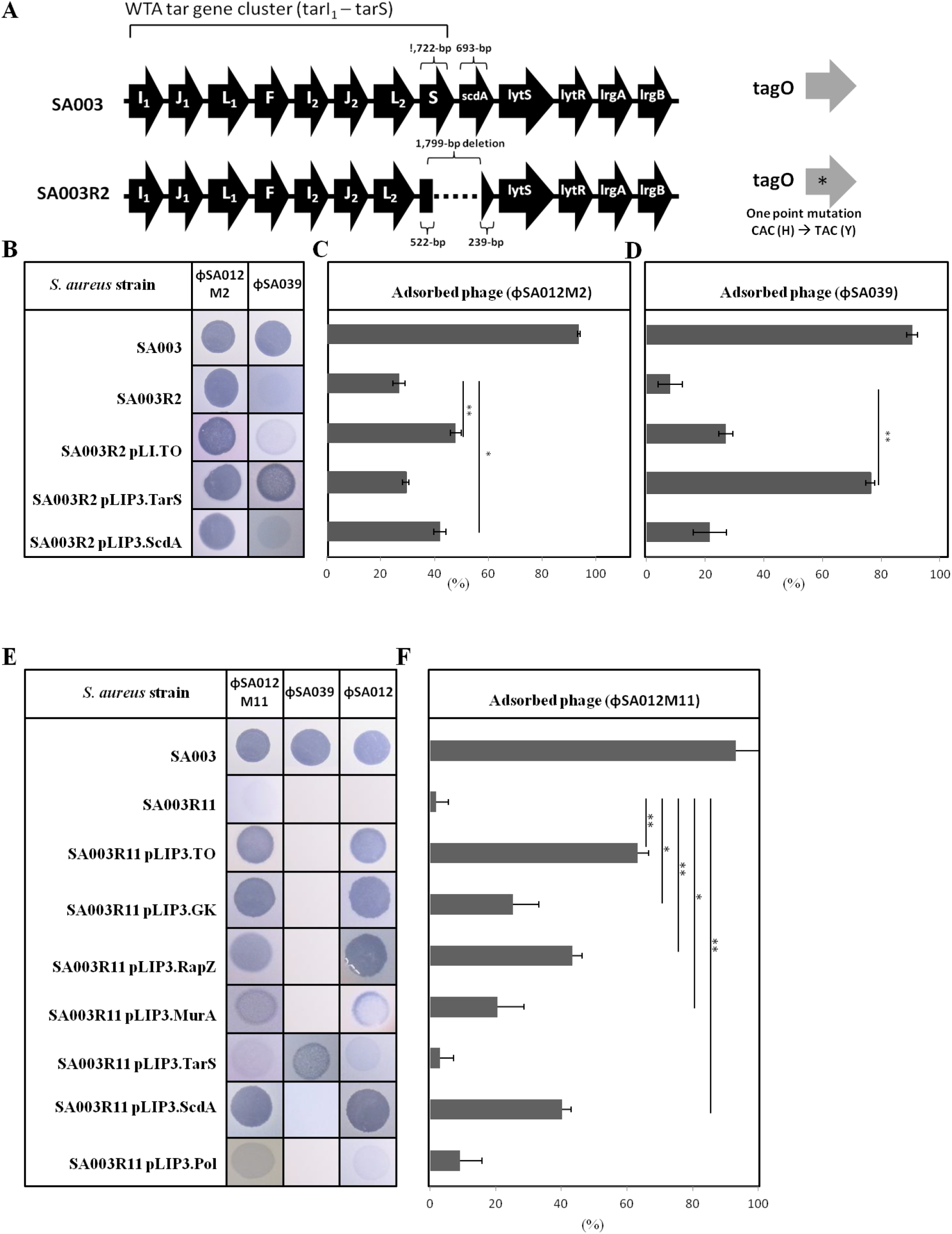
**(A)** Spontaneous mutation in SA003R2; deletion of *tarS* and *scdA* gene, one point mutation in *tagO*. **(B)** Spot test of ϕSA012M2 and ϕSA039 toward SA003R2 and its phage resistance derivative. Adsorption assay of ϕSA012M2 **(C)** and ϕSA039 **(D)** toward SA003R2 and its phage resistance derivative. Spot test **(E)** and adsorption assay **(F)** of ϕSA012M11 toward SA003R11 and its phage resistance derivative. Statistical significance was indicated by *(P<0.05) or **(P<0.01).

Four more genes, *S2121, S768, S515*, and *S2190*, were mutated in the resistant strain, SA003R11, isolated from the 11^th^ round. *S2121*, which encodes UDP-N-acetylglucosamine 1-carboxyvinyltransferase-2 MurA2 (an enzyme which catalyzes the first step of peptidoglycan synthesis together with MurA) showed a nonsense mutation in a glycine-encoding codon (GGA→TGA). *S768*, which encodes a guanylate kinase, and *S2190*, which encodes the alpha subunit of DNA-RNA polymerase, showed point mutations that changed a threonine into isoleucine (ACA→ATA) and an alanine into valine (GCA → GUA), respectively. Mutations in *S156*, *S157*, *S497*, *S2121*, *S768*, and *S2190* were preserved until later generations, including the 20^th^ and 38^th^ rounds, from which we isolated SA003R20 and SA003R38, respectively. In SA003R11 we also identified one point mutation in *S515*, which encodes for the RNAse adaptor protein RapZ. The mutation in this gene developed into two point mutations in SA003R20 and SA003R38. In addition, a point mutation (AAT → GAT) was identified in *S692*, which encodes the putative membrane protein YozB, in SA003R20 and SA003R38.

To evaluate the role of mutated host genes in phage resistance, we performed complementation in SA003R2 and SA003R11 by using wild-type allele in *trans* with each mutated gene. To do so, we employed the *E. coli*/SA shuttle vector pLI50 (Addgene) under promoter P3(Lee et al. 1991; Jeong et al. 2011). As we had identified a nonsense mutation in *S2121*, we also constructed a deletion mutant of this gene. The *S2121* deletion mutant was unchanged in its sensitivity towards ϕSA012 (data not shown).

In resistant strain SA003R2, among three mutated genes *(S156, S157*, and *S497)*, only complementation of *S157* and *S497* can restore phage susceptibility of SA003R2 to ϕSA012M2 (**Fig. 1B**). However, another *Myoviridae* Twort-like phage, ϕSA039, showed completely different infectivity towards SA003R2 and its complemented mutant. ϕSA039 formed faint plaques on SA003R2 plates (**Fig. 1B**), and surprisingly, complementation of *S156* allele in SA003R2 was able to restore the susceptibility to ϕSA039 (**Fig. 1B and 1D**). We suggested that ϕSA039 and ϕSA012 recognize *S. aureus* cell wall by different mechanism. The mutation of SA003R2 is associated with the reduction in WTA production as total phosphate in extracted WTA from SA003R2 decreased significantly compared to wild-type SA003 (**Fig. 2**). WTA-PAGE also showed an invisible ladder of WTA from SA003R2 and all subsequent resistant strains (data not shown).

**Figure 2.**
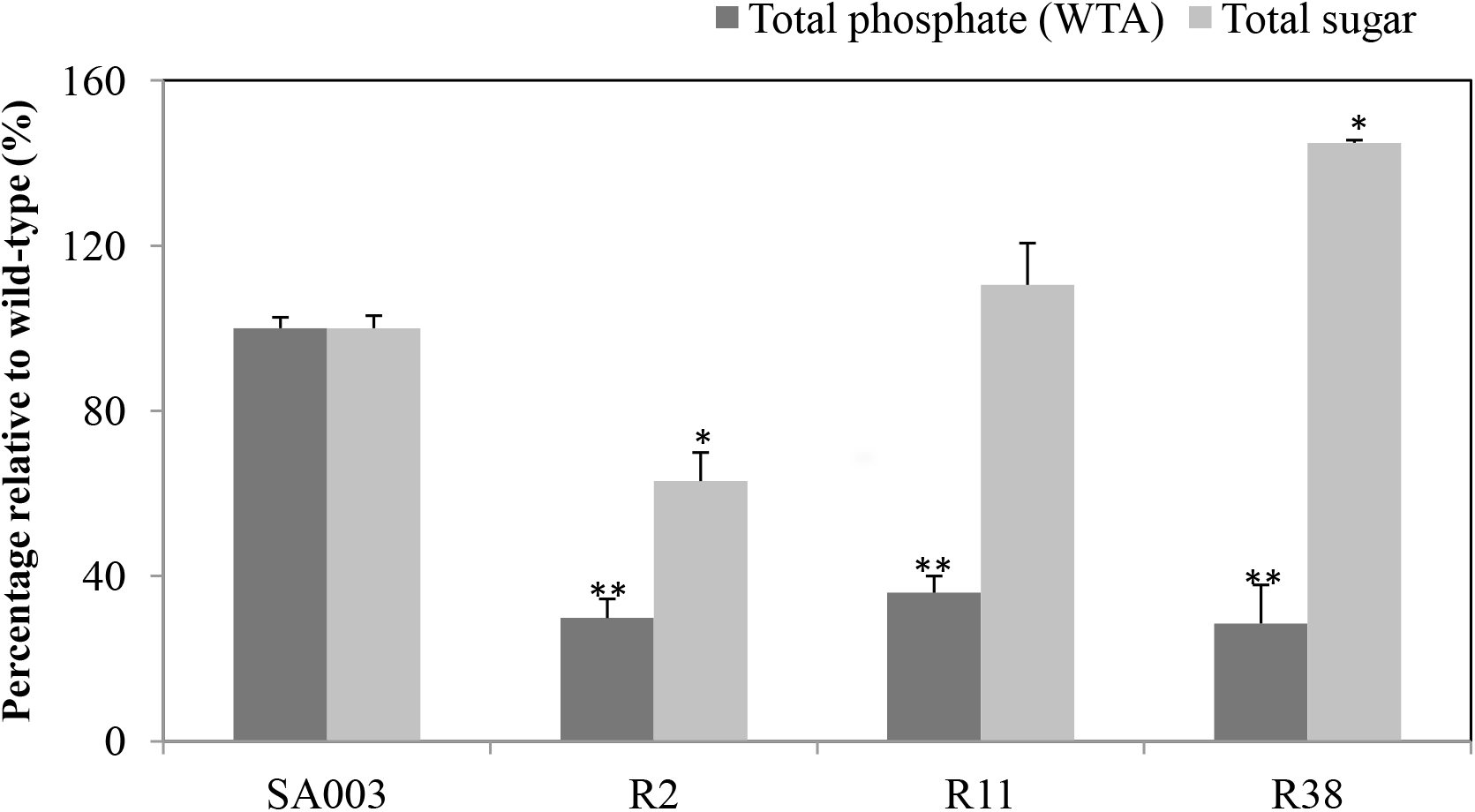
Percentage of total phosphate from extracted wall teichoic acid (WTA) (dark) and whole-cell sugar (gray) relative to the wild-type strain SA003. Values are given as means ± standard deviations (SD, n = 3). Statistical significance was indicated by *(P<0.05) or **(P<0.01).

In resistant strain SA003R11, complementation of each mutated genes, except *S156*, can significantly restore phage susceptibility toward ϕSA012M11 (**Fig. 1E**). The complemented mutant of *S156* has EOP value 5.53±2.52, while complementation of other gene: *S157, S497, S515, S768*, *S2121*, and *S2190*, show EOP value 63.70±1.15, 77.64±0.58, 65.38±0.58, 78.85±0.58, 63.70±1.15, and 43.12±0.58 respectively. Complementation of *S768* and *S497* shows the highest EOP value among other genes. Excluding the *S156* complemented mutant, ϕSA039 failed to form plaques on plates of SA003R11 or its complemented mutants. This observation provides stronger evidence on the importance of *S156* gene for the infection of ϕSA039 as the complementation of other mutated genes did not significantly restore host susceptibility toward ϕSA039.

### Adsorption of ϕSA012 and ϕSA039 to complemented mutant

To determine the adsorption of phages to the complemented mutants, we conducted an adsorption assay for phage ϕSA012M2 and ϕSA012M11 towards wild-type SA003, their phage-resistant counterpart, as well as the complemented mutant. For the complementation in SA003R2 (**Fig. 1C**), adsorption of ϕSA012M2 onto wild-type SA003 and SA003R2 was 93.64±0.44% and 26.83±2.29%, respectively. Complementation of *S156* (R2pLIP3.TarS) did not significantly restore the phage adsorption (29.35±1.15%), while *S497* (R2pLIP3.TO) and *S157* (R2pLIP3.ScdA) restored the phage adsorption to 47.86±1.97% and 42.00±2.30%, respectively. Interestingly, adsorption of ϕSA039 was significantly restored to 76.20±1.49% in R2pLI.TarS indicating the importance of the *S156* gene for the adsorption of ϕSA039 onto SA003.

Next we analyzed the adsorption of ϕSA012M11 toward the complemented mutants of SA003R11. The complemented mutants of SA003R11 harbor *S156* (R11pLIP3.TarS), *S157* (R11pLIP3.ScdA), *S497* (R11pLIP3.TO), *S515* (R11pLIP3.RapZ), *S768* (R11pLIP3.GK), *S2121* (R11pLIP3.MurA2), and *S2190* (R11pLIP3.Pol). As shown in **Fig. 1F**, after 25 min of co-incubation, 93.23±8.79% of ϕSA012M11 was adsorbed onto wild-type SA003 while only 1.92±3.59% was adsorbed onto SA003R11. After complementation of the mutated genes by their respective wild-type alleles, the adsorption of ϕSA012M11 to the complemented mutants of *S157*, *S497*, *S515*, *S768*, and *S2121* was restored. However, even though we observed plaque, the adsorption onto *S2190* complemented mutant did not change significantly. Interestingly, In the case of *S768* gene, even though the EOP value of *S768* complemented mutant (R11pLIP3.GK) was the highest among other genes, the adsorption of phage onto this complemented mutant was relatively low compare to complemented mutant of *S497*, *S515*, and S2121. We speculated that the mutation in *S2190* (which encodes DNA-RNA polymeras alpha subunit), and *S768* (which encodes guanylate kinase) contribute to the inhibition of post-adsorption.

Adsorption of ϕSA012M11 to R11pLIP3.TO, R11pLIP3.ScdA, R11pLIP3.GK, R11pLIP3.RapZ, and R11pLIP3.MurA was restored to 63.20±3.30, 40.25±2.77%, 25.17±7.88%, 43.44±2.95%, and 20.55±7.96%, respectively. Mutation in SA003R11 was associated with an increase in whole-cell sugar according to total carbohydrate quantification using Anthrone reagent (**Fig. 2**). Relative to SA003, the whole-cell sugar of SA003R2 decreased to 62.98±6.92%. Meanwhile, the total sugar of SA003R11 and SA003R38 increased to 110.49±10.10% and 144.89±0.63% relative to SA003, respectively. A slimy texture was visible on the overnight liquid culture of SA003R38 (data not shown).

The infection of wild-type ϕSA012 to the complemented mutant of R11 is similar to ϕSA012M11 according to spot test (**Fig. 1E**). The adsorption of ϕSA012 towards the SA003R11 complemented mutant was not significantly different compared to ϕSA012M11 (**Supplemental Table S1**). However, in most of the genes (except R11pLIP3.ScdA and R11pLIP3.Pol), the wild-type phage’s adsorption showed a tendency to be stronger than ϕSA012M11.

### Peptidoglycan from *S. aureus* inhibits infection of ϕSA012 but not ϕSA039

As some mutated genes (e.g., *S2121* and *S515*) are related to peptidoglycan synthesis, we assume that peptidoglycan may play an important role in inhibiting phage adsorption. For this reason, we performed a plating efficiency test with the addition of peptidoglycan. Different concentrations of peptidoglycan from *S. aureus* (Invivogen) were added to 100 μl of 10^3^ PFU/ml phage prior to the plaque assay against SA003. Two different phages, ϕSA012 and ϕSA039, belonging to the same group of *Myoviridae* Twort-like phages, were used. We used the *E. coli* phage SP-21(Tanji et al. 2005) as a negative control. As shown in Fig. 3, peptidoglycan inhibited the infection of ϕSA012 but not ϕSA039. To confirm if O-acetylation in C6 of the muramic acid residue is important for binding of ϕSA012, we deleted *oatA* gene by CRISPR/Cas9 system using pCasSA plasmid. However, the adsorption of ϕSA012 onto *oatA* deletion mutant did not change (**Fig. 5**).

**Figure 3.**
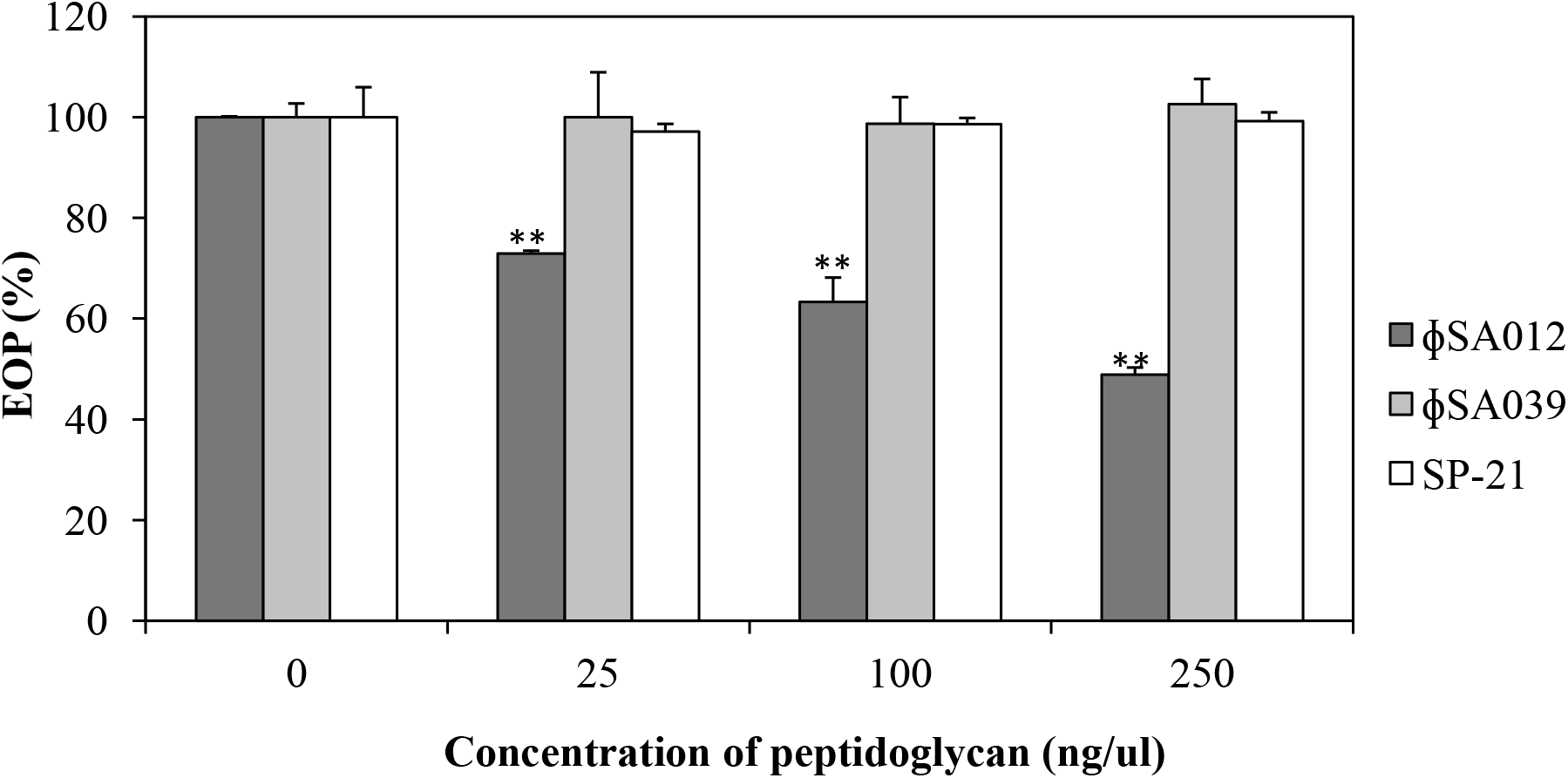
Dose-dependent inhibition by peptidoglycan of phage ϕSA012 (black bar), phage ϕSA039 (gray bar) and *E. coli* phage SP-21 (white bar). Data represent means ± standard deviations (SD, n = 3). Statistical significance was indicated by *(P<0.05) or **(P<0.01).

### Genome comparison of ϕSA012 and ϕSA039

According to morphological observation under TEM, ϕSA039 and ϕSA012 belong to *Myoviridae* phages. Both of them show broad host ranges against *S. aureus* isolated from the milk of mastitic cows (Synnott et al. 2009). As these two phages show different infectivities, we analyzed the whole genome of ϕSA039 and mapped it to the genome of ϕSA012 as a reference. Whole genome of ϕSA012 has been explained in previous report (Takeuchi et al. 2016). Whole genome sequencing of ϕSA039 revealed that its genome size is 141,038 bp long and contains 217 ORFs. The ϕSA039 genome encodes three tRNAs (pseudo Trp-tRNA gene, bp 27,174 to 27,103; Phe-tRNA gene, bp 27,253 to 27,181; and Asp-tRNA gene, bp 27,333 to 27,260). At the nucleotide level, ϕSA039 and ϕSA012 share 96% similarity where several genes are missing or partially deleted in either ϕSA012 or ϕSA039 (listed in **Supplemental Table S2**). The genome of ϕSA039 shares 99% similarity with staphylococcal phage JD007 (Cui et al. 2012), which is classified as a member of the *Myoviridae* phages that are serotype D *S. aureus*–specific lytic phages that include the well-known phage K (O’Flaherty et al. 2004).

As ϕSA039 and ϕSA012 differ in their host preference, we select the region of genome encoding the putative tail and baseplate protein and aligned the open reading frames (ORFs) in that region for amino acid alignment by basic local alignment tool (blastp). Most of the ORFs in the selected region share a high degree of similarity (**Fig. 4**); most of the ORFs have more than 99% similarity and three ORFs share 91% similarity. Three ORFs share relatively low similarity with all the others: *orf100* (86%), *orf101* (88%), and *orf96* (83%). Moreover, *orf96* of ϕSA039 lacks 195 amino acid residues compared to ϕSA012. *Orf100* and *orf102* of ϕSA039 are homolog of *orf103* and *orf105 of* ϕSA012 which were previously reported as receptor-binding protein (RBP).

**Figure 4.**
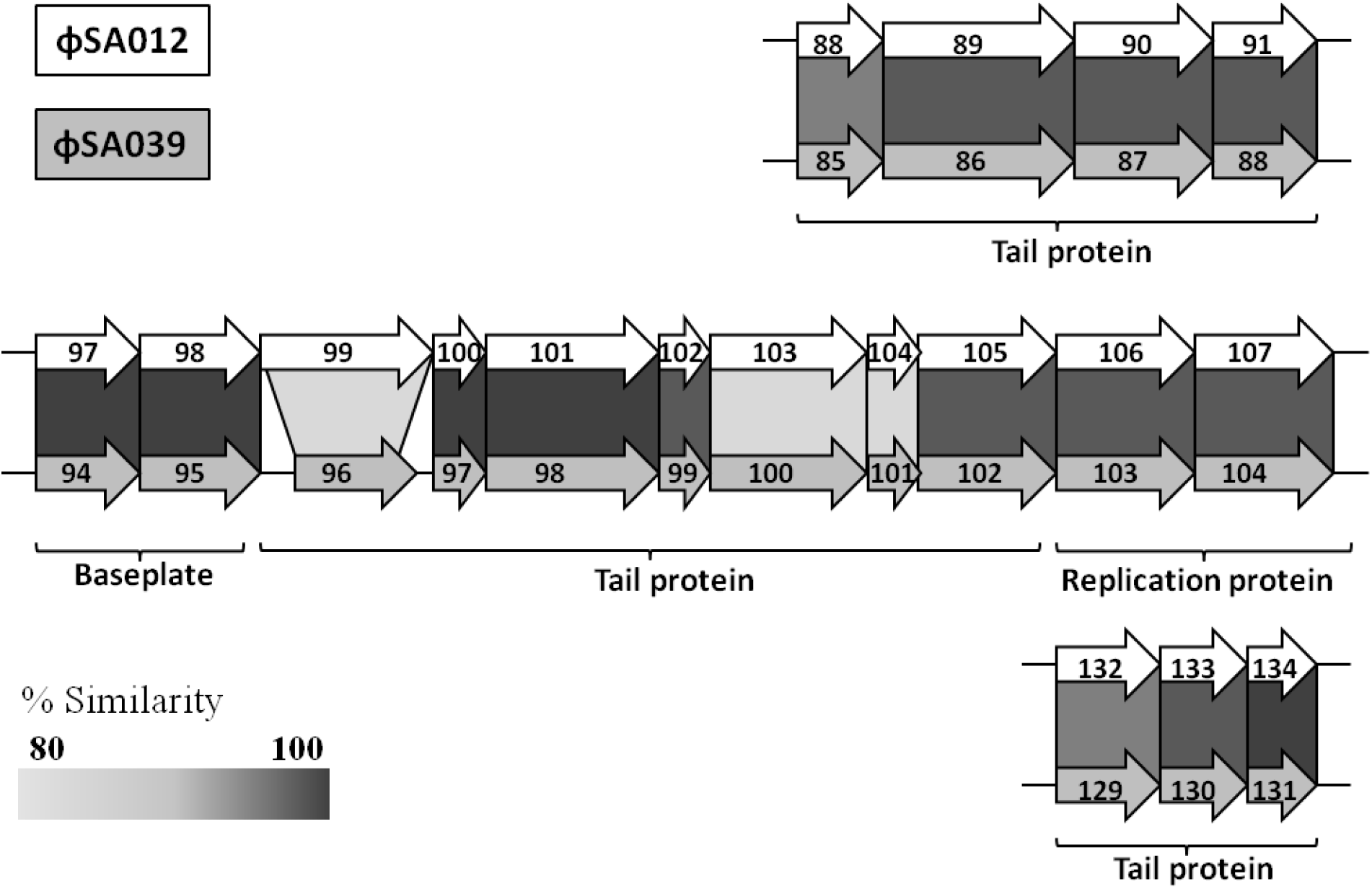
Alignment of putative tail and baseplate region between ϕSA012 and ϕSA039.

To determine whether protein of *orf100* and *orf 102* of ϕSA039 has same activity with their homolog genes, we performed neutralization test in ϕSA039 by EOP with antibodies against *orf103* protein of ϕSA012 (anti-ORF103 serum) and *orf105* protein of ϕSA012 (anti-ORF105). Infection in SA003 by ϕSA039 was completely inhibited by anti-ORF105 serum but not by anti-ORF103 serum (data not shown).

### Artificial deletion of *tarS* gene in SA003 significantly reduce the adsorption of ϕSA039 but not ϕSA012

WTA has been well reported as a phage receptor in *S. aureus* (Xia et al. 2011). Phage K belongs to the same group as ϕSA012 and ϕSA039 is able to bind to the backbone of WTA. Phage K and the other *S. aureus* phages lost their infectivity towards the *tagO* deletion mutant. As described earlier, phage susceptibility of SA003R2 toward ϕSA039 was decreased drastically, and complementation of *S156* gene for glycosyl transferase TarS, enzyme that glycosylate β-GlcNAc on WTA, restore the adsorption of ϕSA039. We hypothesized that β-GlcNAc on WTA is crucial for binding of ϕSA039. To test our hypothesis, we first generated a deletion and complementation of *tagO* in *S. aureus*. As shown in **Fig. 5**, our spot test and adsorption assay failed to detect infection of the *tagO* deletion mutant (RNΔTagO) either by ϕSA039 or ϕSA012. Complementation of the *tagO* deletion (RN△TagO::plIP3.TO) restored the adsorption of ϕSA039 and ϕSA012 to 50.55±2.50% and 57.75±1.57%, respectively. On the other hand, deleting *tarS* caused the significant reduction only on the adsorption of ϕSA039. Complementation of *tarS* deletion mutant (SA003△TarS::pllP3.TarS) restored the adsorption of ϕSA039 to 88.69±0.98%. This result indicated that ϕSA039 requires β-GlcNAc on WTA. Since the adsorption of ϕSA039 was not completely loss in TarS null SA003, this phage may also use backbone of WTA for its binding.

**Figure 5.**
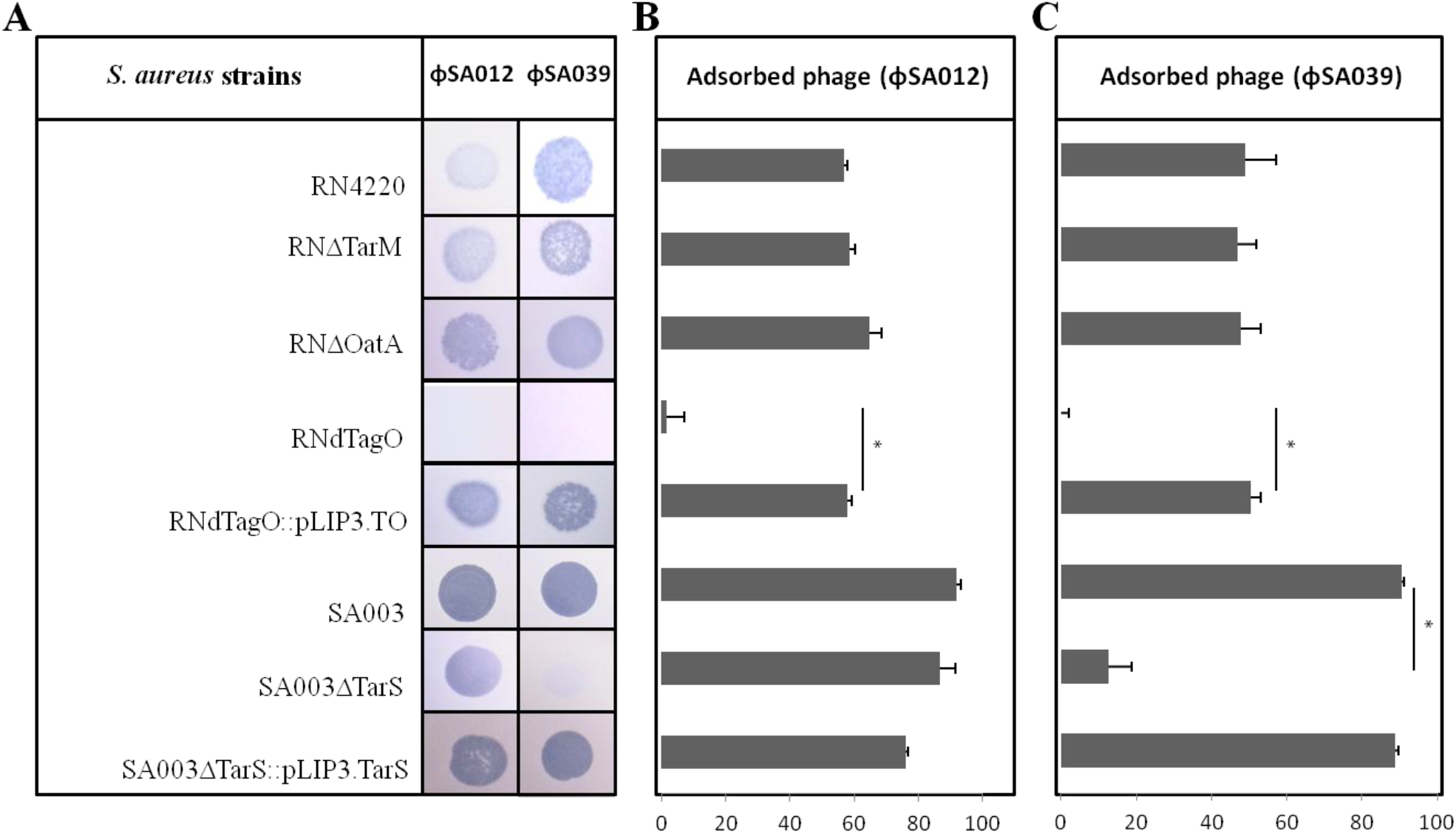
Infectivity of ϕSA012 and ϕSA039 towards RN4220, *tarM* deletion mutant (RN△TarM), oatA-deletion mutant (RN ΔOatA), tagO-deletion mutant (RNdTagO), and complemented *tagO* (RNdTagO::pLIP3.TO). tarS-deletion mutant (SA003ΔTarS) and complemented *tarS* (SA003ΔTarS::pLIP3.TarS) The infectivity assay is based on spot test (A) and adsorption assay for ϕSA012 (B) and ϕSA039 (C). Data represent means ± standard deviations (SD, n = 3). Statistical significance was indicated by *(P<0.05) or **(P<0.01).

## Discussion

Our results show that genotypic changes occurred during coevolution of bacteria and phage. Most of the mutated genes in phage resistant SA003 were linked to phage adsorption inhibition. In the early round of co-culture, SA003R2 altered its WTA which is well known as a phage receptor in *S. aureus*. To counter the mutant host, ϕSA012 phage altered its Receptor-binding protein (RBP) (Takeuchi et al. 2016). However, in later round, in response to the counteradaption of phage, coevolved SA003 from 11^th^ round (SA003R11) changed its cell wall structure, overproduced the capsular polysaccharides, and inhibited the post-adsorption step by acquiring a point mutation in the DNA/RNA polymerase alpha subunit and guanylate kinase. Finally, the later coevolved SA003R38 acquired a point mutation in another gene encoding putative membrane protein which seems to be involved in the excessive production of polysaccharide.

### Most mutated host genes are linked to phage adsorption

In the early round of coevolution, phage resistant strain SA003R2 significantly reduces the adsorption of ϕSA012. Wall Teichoic Acid (WTA) has been previously characterized as a phage receptor for *S. aureus*. Consistent with these observations, three mutated genes *(S497, S156*, and *S157*) are associated with WTA. *S497* (TagO) is involved in the initial synthesis of WTA particularly in the attachment of WTA to peptidoglycan (Soldo et al. 2002), *S156* (TarS) glycosilates β-GlcNAc on WTA (Brown et al. 2012), and *S157* (ScdA) affect cell wall morphology. Deletion in *scdA* has been reported to increase the peptidoglycan cross-linking (Brunskil et al. 1997). However, as the complementation of *S156* did not significantly restore susceptibility of SA003R2 to ϕSA012, we assumed that ϕSA012 does not require β-GlcNAc moieties in WTA. The mutation in SA003R2 was followed by a decrease in total phosphate from extracted WTA, which may indicate the alteration of WTA production. However, ϕSA012 has been shown to have at least two RBPs (ORF103 and ORF105) (Takeuchi et al. 2016). Therefore, the phage may alter one of them to enhance the binding ability to its host. The mutant phage counterpart of SA003R2, ϕSA012M2, harbors two point mutations in *orf103* consistent with the increase in infectivity towards SA003R2. We concluded that ϕSA012M2 developed mutation in *orf103* to fasten the binding onto SA003R2.

In response to the evolution of ϕSA012M2, SA003R11 acquired point mutations in four other genes: *S515* (*rapZ*), *S768* (Guanylate kinase), *S2121* (*murA*), and *S2190* (DNA-RNA polymerase alpha subunit). Three of them (*S2121, S515*, and *S768*) are associated with phage adsorption. MurA2 (*S2121*) has been reported to catalyze the first step of peptidoglycan synthesis together with *murA* (Blake et al. 2009). A nonsense mutant of this gene may alter peptidoglycan synthesis and inhibit phage adsorption to *S. aureus* either directly or indirectly. However, our study shows that deletion mutant of *murA2* did not change their susceptibility toward our tested phage. It is likely that the effect of mutation in *MurA2* gene is related to another mutated gene.

RapZ (S515) modulates the expression of GlmS which was reported to be the key enzyme that feeds glucose into cell wall synthesis in *S. aureus* (Komatsuzawa et al. 2004). It is indirectly involved in the assembly of cell wall components such as peptidoglycan, and either lipopolysaccharide in Gram-negative bacteria or lipoteichoic acid in Gram-positive bacteria. It is also involved in capsule or exopolysaccharide production (Coley and Baddiley 1972; Plumbridge et al. 1993). As whole-cell sugar was higher in SA003R11 (which harbors a point mutation in *S515*) than in SA003R2, the host’s cell wall may change due to either overproduction of capsular polysaccharide or alteration of peptidoglycan. Moreover, SA003R38, which harbors two point mutations in RapZ (*S515*), had much higher whole-cell sugar than SA003R11. Mutations in *S515* are therefore likely to be related to the overproduction of capsular polysaccharide. Our result is consistent with previous observations that a capsule inhibits bacteriophage infection (Bernheimer and Tiraby 1976; Scholl et al. 2005). Our finding provides deeper insight on the analysis of the *rapZ* gene involved in the capsular polysaccharide production of coevolved *S. aureus* in their defense against phage infection. In addition, in the later round (SA003R38), a point mutation was found in *S692* encoding putative membrane protein YozB. As slimy texture and significant increase of whole-sugar were observed in SA003R38, gene *S692* may also be involved in capsule production.

Guanylate kinase (*S768*) is involved in a crucial intermediate step in RNA/DNA synthesis by catalyzing guanosinemonophosphate (GMP) phosphorylation to form guanosine diphosphate (GDP) (Konrad 1992). This enzyme is broadly distributed nucleotide kinase and essential for cellular GMP recycling and nucleotide equilibrium (Oeschger 1978). This enzyme participates in purine metabolism (Weber et al. 1992). In other studies, purine biosynthesis has been shown to be associated with the survival of *S. aureus* under conditions of stress, such as the presence of vancomycin and daptomycin (Keer et al. 2001; Mongodin et al. 2003). Yee et al. (2015) reported that a defect in purine biosynthesis pathway may be related to downstream energy production (i.e., increased purine biosynthesis fuels the generation of polymers, the most energy-demanding metabolic process in bacteria). Two of the most abundant polymers in *S. aureus* and *B. subtilis* are peptidoglycan (Mongodin et al. 2003) and WTA (Ellwood 1970). Our study shows that the complement mutant of this gene show high EOP value compared to other complement mutant but relatively low adsorption rate. We interpreted that the mutation of this gene is related to both inhibition of phage adsorption and post-adsorption. By affecting purin metabolism, the mutation is likely involved in the recycling of injected phage-DNA, and subsequently fueling the generation of polymers in the cell wall such as teichoic acid and capsular polysaccharide.

Complementation in *S2190* (DNA–RNA polymerase) did not restore phage adsorption but did increase the EOP value. Mutations in *S2190* may not be associated with phage adsorption, but with post-adsorption. RNA polymerase alpha subunit has been reported to be involved in transcription activation (Ishihama 1992). Most phages with a genome size less than 200 kbp commonly did not harbor a complete set of genes responsible for genome replication and nucleotide metabolism. Only large-genome size phage (termed Jumbo phage), with genome larger than 200 kbp, have been reported to have more than one paralogous gene for DNA polymerase and RNA polymerase (RNAP) (Yuan and Gao 2017). Our phage in this study has a genome size around 142 kbp and the complete set paralogous of RNA polymerase subunit was not found (Takeuchi et al. 2016). Therefore this phage may use host RNA polymerase as machinery for their DNA transcription. Hence, the mutation that we observed in this gene is likely to be involved in the inhibition of phage DNA transcription. This finding is supported by a previous observation (Osada et al. 2017) that SA003R11 resists phage infection not only by inhibiting phage adsorption but also by suppressing phage genome replication.

In this study, we also observed that the infection of wild-type ϕSA012 onto the complemented mutant of R11 was not significantly different compared to ϕSA012M11. However, excluding R11pLIP3.ScdA and R11pLIP3.Pol, the adsorption of wild-type ϕSA012 showed a tendency to be stronger than ϕSA012M11. Fitness cost might be detected in ϕSA012M11 as it experienced long term coexistence with SA003. Therefore, the infection of ϕSA012M11 toward the R11 complemented mutants is weaker than that of wild-type ϕSA012. Another report has also demonstrated a fitness cost for phage (which experienced coevolution) by the reduction of its infectivity toward wild-type host, or by the limitation of host range expansion (Hall et al. 2011).

CRISPR/cas motif was not found in the genome of wild-type SA003 and short fragments of phage DNA of ϕSA012 were not found in genome of phage resistance derivative, which indicates that the bacteria likely did not use CRISPR/cas to defense the phage invasion. It is well known that in many pathogenic bacteria, including *S. aureus*, many of the clones have lost CRISPR/cas during evolution. As a result, *S. aureus* frequently exchange their genetic material via phage-mediated horizontal gene transfer (Brussow et al. 2004; Lindsay 2010). Our study provides deeper insights on how phage and its host can co-exist during co-culture and will help guide future assessments of the advantages and disadvantages of phage therapy.

### Two *Myoviridae* Twort-like phages (ϕSA012 and ϕSA039) use different mechanisms to infect *S. aureus*

*Myoviridae* Twort-like phages are well known for their wide host range and highly lytic capabilities. Phages belonging to this genus have been considered promising candidates for therapeutic use and as detection agents of bacterial contamination (Loessner et al. 1996). Recently, Alves et al. (2014) showed that Twort-like phages were effective at reducing biofilm formation by *S. aureus* strains. Our data showed that although the ϕSA012 and ϕSA039 genomes share high similarity, these two phages infect *S. aureus* in different ways. ϕSA039 infectivity towards SA003R11 and SA003R2 and its complemented mutant is lost or significantly lessened, except in the instance of *S156* complementation. As a result, we conclude that ϕSA039 use *S156* for its binding to *S. aureus* cell.

Comparison of ϕSA012 and ϕSA039 binding mechanism was performed in this study. First, we found that both phages bind to WTA. We verified that the deletion mutant of *tagO* is resistant to both phages. Second, we found that peptidoglycan from *S. aureus* was able to block infection of ϕSA012 but not ϕSA039. It is reported that peptidoglycan can block infection of *Siphoviridae* phages Coyette and Ghuysen (1968). Li et al. (2016) determined that ϕ11, one of the best-studied *Siphovirus* es, requires O-acetylated peptidoglycan for adsorption. However, our data in this study suggests that ϕSA012 does not require O-acetylated peptidoglycan for adsorption (**Fig. 5B**). We assumed that ϕSA012 uses another component in the peptidoglycan polymer. Finally, we confirm that *S156* which encodes β-N-acetyl glucosaminidase TarS, is crucial for ϕSA039. This suggests that β-N-acetyl glucosamine in WTA acts as a receptor of ϕSA039. Most *S. aureus* synthesize repeating unit of ribitol-phosphate (RboP)WTA with three tailoring modification D-alanine, α-GlcNAc, and β-GlcNAc (Brown et al. 2013). The GlcNAc moieties are attached to RboP by two independent enzymes namely α-GlcNAc WTA glycosyltransferase TarM (Xia et al. 2010), and β-GlcNAc WTA glycosyltransferase TarS (Brown et al. 2012).

Most *S. aureus* phages target WTA and its GlcNAc moieties for adsorption. Notably, it has been reported that *Siphoviridae* phages use α-GlcNAc moieties as receptors (Xia et al. 2011; Li et al. 2016), while *Podoviridae* phages, such as ϕ44AHJD, ϕ66, and ϕP68, use β-GlcNAc moieties (Li et al. 2015). In contrast, *Myoviridae* phages have been reported to simply require WTA polymer, regardless of GlcNAc acetylation (Xia et al. 2011). It is not widely known that phages belonging to the *Myoviridae* group also require GlcNAc moieties in WTA. Our study shows that ϕSA039, a *Myoviridae* phage, with 96% similarity to phage ϕK, a well known *Myoviridae* phage, requires β-GlcNAc in WTA for its binding. In addition, we also confirmed that the *tarM*deletion mutant and its wild-type *S. aureus* have the same susceptibility to ϕSA039 (**Fig.5**). Hence, α-N-acetyl glucosamine of WTA is likely not essential for ϕSA039. As β-N-acetyl glucosamine has been reported to serve as a binding site of PBP2a, the enzyme responsible for β-lactam antibiotic resistance in *S. aureus* (Brown et al. 2012), the application of ϕSA039 and β-lactam antibiotic may give a synergetic effect for the treatment of *S. aureus* infection. We believe that this finding may give us novel insights into how closely related phages in the group of *Myoviridae* differ in their host preference among *S. aureus* strains.

### *In silico* analysis of ϕSA039 and ϕSA012 genome shows potential viral protein that contributes to different adsorption mechanism

Several genes are missing in ϕSA012 and ϕSA039 (**Supplemental table S2**). In particular, two terminal repeat regions (*orf195* and *orf200*) that exist in ϕSA012 genome are missing in ϕSA039. Lobocka et al. (2012) reported that all genes located in the terminal redundant region play a role in host takeover that is analogous to *B. subtilis* phage, SPO1, which also possesses a terminal redundant region (Stewart et al. 1998). In addition, the gene of *orf81* of ϕSA012, which contains a putative intron-encoded nuclease, and the gene of *orf39* of ϕSA012, which encodes a DNA ligase are also missing in ϕSA039. Moreover, a partial deletion is found in *orf96* of ϕSA039, a homolog of *orf99* of ϕSA012, which encodes tail morphogenetic protein. As we described in a previous study, the mutant phage ϕSA012M20 carries a point mutation in *orf99*, and subsequently accumulates two more point mutations to become the mutant phage ϕSA012M38 (Takeuchi et al. 2016), which may indicate the importance of *orf99* in phage-host interaction. Therefore, the different in *orf96* of ϕSA039 may contribute to the specificity different.

In the tail or baseplate region of ϕSA039, compared to ϕSA012, several ORFs have been shown to have low amino acid similarity (83–88%): i.e., *orf99, orf103*, and *orf104*, whereas *orf103* and *orf104* are located in the unique region of the Twort-like tail/baseplate module (**Fig. 4**). During phage-host interaction, *orf103* of ϕSA012 had been reported to bind onto α-GlcNAc of WTA, while *orf105* is likely bind onto the backbone of WTA. Since anti-ORF105 of ϕSA012 can neutralize of ϕSA039 infection, the binding activity of *orf102* of ϕSA039, a homolog of *orf105* of ϕSA012, must be similar. Thus, in ϕSA039, *orf102* may bind onto backbone of WTA. However, *orf100*, a homolog of *orf103* of ϕSA012, may bind to β-GlcNAc or another part of WTA. Taken together, we suspect that ORFs of ϕSA039 located in the tail or baseplate region, which has low amino acid similarity compared to ϕSA012 such as *orf100, orf101*, and *orf96*, are likely to be the factors responsible for the observed differences in specificity between ϕSA012 and ϕSA039. However, further analysis of those potential genes might be necessary.

## Acknowledgments

We would like to thank Professor Takehiko Itoh (School of Life Science and Technology, Tokyo Institute of Technology) for allowing us to use NGS analysis in his lab. We also thank Professor Masaaki Wachi (School of Life Science and Technology, Tokyo Institute of Technology) for his useful advice.

## Ethical approval

This article does not contain any studies with human participants and animals performed by any of the authors.

